# Clinical evidence yield as a framework for evaluating computational predictors and multiplexed assays of variant effect

**DOI:** 10.64898/2026.03.27.714777

**Authors:** Yifei Shang, Mihaly Badonyi, Joseph A. Marsh

## Abstract

Interpreting the clinical significance of missense variants of uncertain significance (VUS) remains a major challenge in clinical genetics. Although computational variant effect predictors (VEPs) and multiplexed assays of variant effect (MAVEs) can generate large-scale functional scores, their value is typically assessed using discrimination metrics such as AUROC rather than by the strength of evidence they provide under ACMG/AMP guidelines. Here, we introduce mean evidence strength (MES), a quantitative metric that summarises the pathogenic and benign evidence assigned across missense variants following gene-level Bayesian calibration. Using the acmgscaler framework, we calibrated 12 population-free VEPs across 367 disease genes and analysed 15 MAVE datasets with sufficient clinical data. MES revealed important discrepancies with AUROC, including cases where methods with similar discrimination differed substantially in evidence yield. MAVEs achieved high average MES despite lower AUROC, while several VEPs showed strong discrimination but more limited calibrated evidence. Among predictors, CPT-1 achieved the highest MES and provided moderate or stronger evidence for the largest fraction of ClinVar VUS. MES therefore provides a practical framework for evaluating computational and experimental variant effect datasets in terms of calibrated clinical evidence yield.

## Introduction

Deciphering the genotype-phenotype relationship is a fundamental goal of genetics. In the clinical context, timely diagnosis and therapeutic interventions in precision medicine critically depend on the accurate mapping from patient mutations to clinical phenotypes. However, although advances in genome sequencing capacity have uncovered millions of human variants, interpreting the clinical implications of these variants remains challenging. As a result, the vast majority of recorded variants remain variants of uncertain significance (VUS) [1]. Among these VUS, missense variants, with the potential to directly alter protein structure and function, represent a particularly large and growing clinical concern [2]. To address this problem, the American College of Medical Genetics and Genomics and the Association of Molecular Pathology (ACMG/AMP) provided guidelines for combining distinct lines of evidence to evaluate variant pathogenicity or benignity [3]. Such evidence supporting or refuting pathogenicity can come from various sources, but the current ACMG/AMP system relies heavily on the retrospective analysis of variant population frequency and segregation data.

Recently, emerging proactive and high-throughput approaches serve as powerful tools to tackle the missense VUS problem. Computational variant effect predictors (VEPs) are machine learning models capable of estimating missense variant pathogenicity at proteome scale [4]. Experimental multiplexed assays of variant effect (MAVEs) interrogate the functional consequences of missense variants in a high-throughput manner [5]. Despite their scalability, the clinical utility of MAVEs and VEPs is often underestimated for two main reasons. First, under current ACMG/AMP guidelines, VEPs are limited to supporting evidence, and while “well-validated functional assays” can in principle provide strong evidence, the criteria for such validation are ambiguous [3]. Second, VEPs and MAVEs are typically evaluated based on their ability to distinguish between known pathogenic and benign missense variants using metrics such as the area under the receiver operating characteristic curve (AUROC). While these measures capture retrospective classification performance on labelled variants, it remains unclear whether they fully reflect the clinical utility of these datasets for variant interpretation.

Recently, several calibration methods compatible with the ACMG/AMP framework have been proposed to convert functional and computational variant effect scores into standardized evidence strengths for clinical variant interpretation. For example, functional assay results can be calibrated by comparing the distributions of known pathogenic and benign variants in functionally normal and abnormal variants to derive odds of pathogenicity values that map directly to standardized ACMG/AMP evidence strengths [6]. However, this approach relies on fixed thresholds that assign the same evidence category to all variants beyond a cutoff, which can inflate evidence for variants close to the boundary.

A more recent method proposed for VEPs estimates the likelihood ratios of pathogenicity from the local density of pathogenic and benign variants across the score distribution, which can be translated into distinct evidence strength categories in ACMG/AMP guidelines [7]. Although this strategy has improved the integration of computational predictions into clinical interpretation [8], genome-wide calibration can produce heterogeneous performance across genes [9]. This has motivated the development of gene-level calibration approaches for both VEP and MAVE datasets.

Multiple approaches for gene-level calibration have recently been proposed. MaveLLR applied gene-level calibration to a MAVE dataset for HMBS [10], while other approaches model MAVE score distributions to estimate gene-specific prior probabilities of pathogenicity [11]. A recent data-adaptive framework for calibrating VEP scores can aggregate variants at the gene or domain level [12]. We developed *acmgscaler*, an R package for gene-level calibration of variant effect scores that estimates likelihood ratios from the distributions of pathogenic and benign variants using bootstrapped kernel density estimation [13].

The advent of these gene-level calibration tools motivated us to quantify how much clinical evidence VEP and MAVE data can provide, and how this compares to traditional performance metrics such as AUROC. The limitations of relying on AUROC alone were highlighted in a recent study of RYR1 variants, where a VEP achieved strong classification performance but generated limited clinical evidence because pathogenic and benign variants showed substantial score overlap [14]. To address this, we define mean evidence strength (MES), a metric that averages the pathogenic and benign point values across the full variant score distribution to quantify the total clinical evidence yield of a dataset. We observed notable discrepancies between classification performance on known labels and clinical evidence yield, indicating that AUROC alone may not capture the clinical utility of VEP and MAVE datasets. This framework provides an interpretable way for VEP and MAVE datasets to be evaluated in terms of their clinical evidence yield for genetic diagnosis.

## Results

### Mean Evidence Strength Quantifies the Clinical Utility of VEP and MAVE Scores

We first used our recently introduced *acmgscaler* tool to perform gene-level calibration of VEPs using pathogenic/likely pathogenic and benign/likely benign missense variants from ClinVar as the truth set. Importantly, we considered only *population-free* VEPs, as defined in our recent study [15], because they have not been trained on, or otherwise exposed to, human clinical or population variants, allowing the full set of known pathogenic and benign variants to be used for calibration without risk of circularity. In contrast, VEPs trained on variants that are subsequently used for calibration may overstate evidence strengths. Even *population-tuned* models, such as AlphaMissense [16], which are not directly trained on clinical labels, incorporate allele frequency information that is often used for clinical variant classification, and are therefore also susceptible to circularity in this setting [17]. However, we do include popEVE [18]; although, strictly speaking, it is exposed to population variation, this is used only for per-gene score adjustment and preserves the within-gene ranking of variants. Because our calibration depends on within-gene rank, this avoids any risk of circularity.

After calibration, each variant effect score is assigned an evidence strength by *acmgscaler*, reflecting the likelihood ratio for pathogenicity under the Bayesian ACMG/AMP framework. These likelihood ratios can then be converted into evidence points using a quantitative scoring, in which each evidence category contributes a defined number of points toward variant classification [19]. We defined mean evidence strength (MES) as the average of the absolute evidence point values assigned to variants in a dataset (Figure 1).

**Figure 1.**
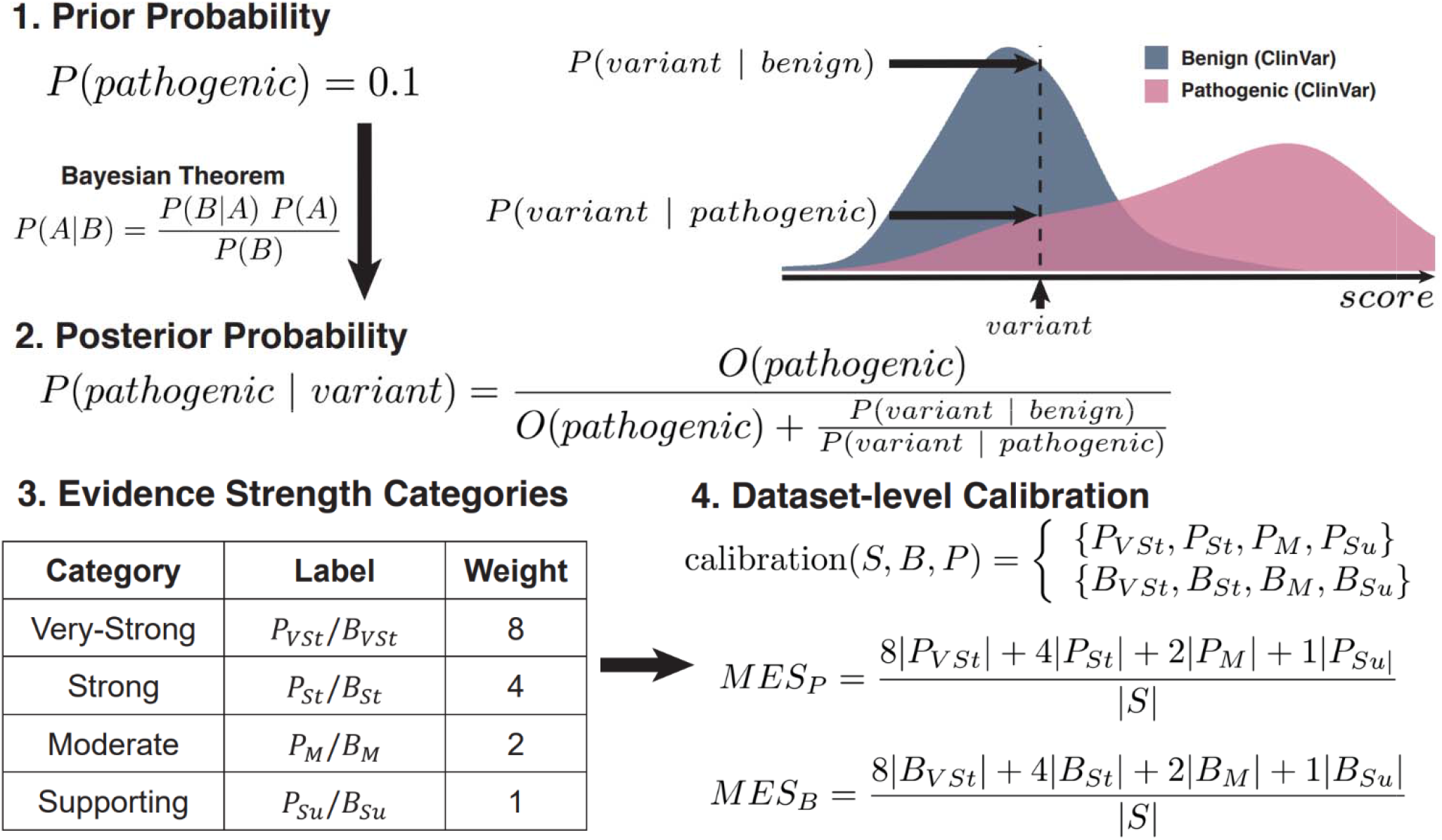
Bayesian variant score calibration and clinical evidence calculation pipeline. Overview of the Bayesian gene-level variant score calibration using known labels and definition of the MES metric. MES is divided into pathogenic and benign components scored based on variant evidence categories after calibration. denotes the set of all input variant scores in a gene. and denote the label sets for benign and pathogenic variants respectively.

Different predictors vary in their coverage across variants: some do not provide scores for all genes, while others lack scores for certain variants within genes. For a fair comparison between VEPs, we therefore restricted the analysis to variants with scores available across all predictors. First, we considered 12 VEPs with relatively high coverage, with scores for 2.05 million variants across 367 human disease genes for which calibration could be performed (note that *acmgscaler* requires a minimum of 10 pathogenic and 10 benign variants per gene for reliable density estimation). In Fig. 2a, we rank these VEPs by their average MES across all genes. Interestingly, CPT-1 [20] ranks first, consistent with our previous study that found it to be the top-performing model in terms of correlation with MAVE data across 97 VEPs [15]. In Fig. S1, we include three additional VEPs with lower coverage, considering scores for 974,071 variants across 200 genes. Here, EVE [21] ranks slightly higher than CPT-1. However, since EVE only provides predictions where sufficient alignment depth is available, its performance may be biased towards high-confidence variants, potentially inflating the observed results.

**Figure 2.**
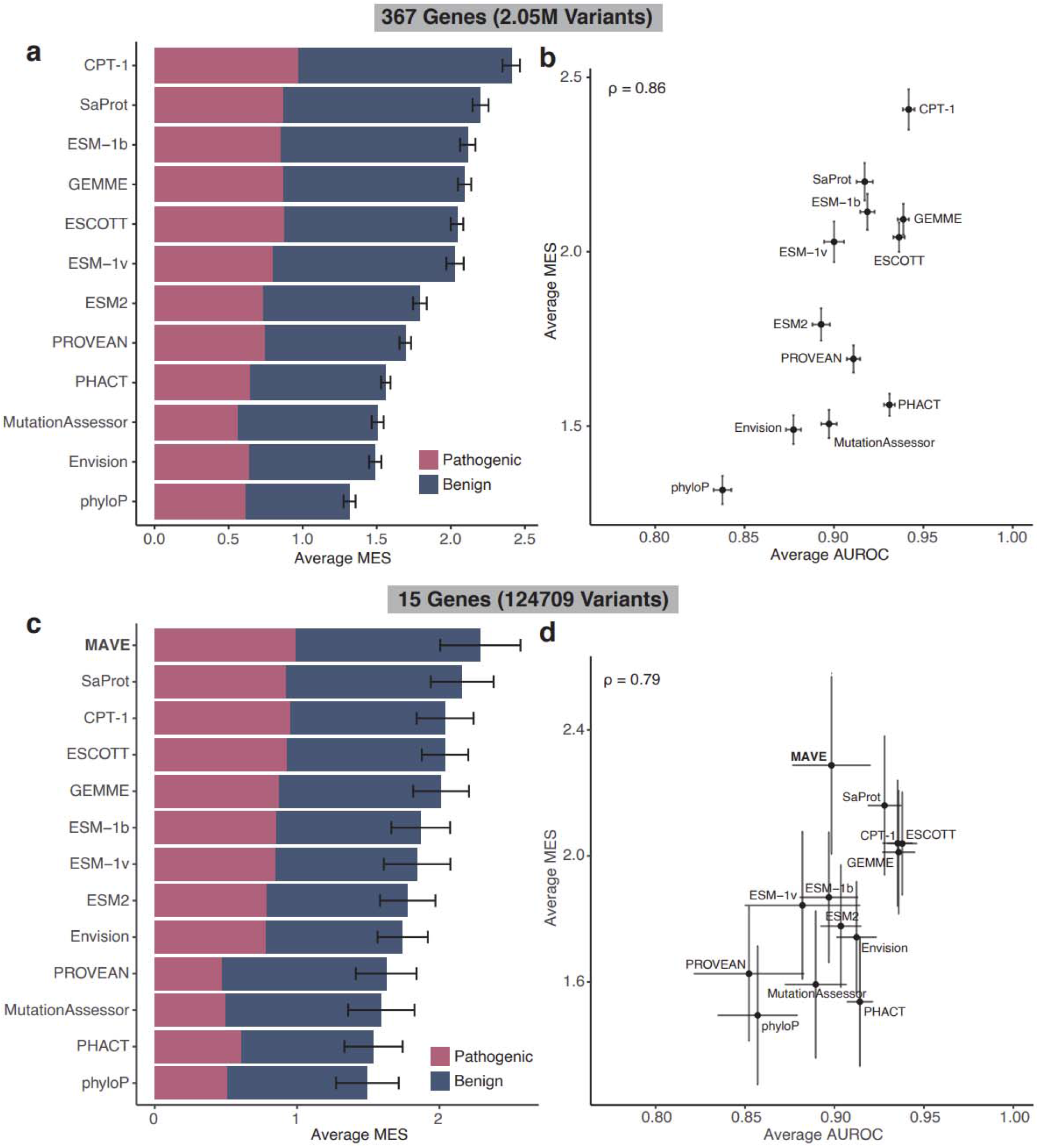
Measuring clinical utility of VEPs and MAVEs using the MES metric. **a,c**. Average MES, split into pathogenic and benign components, when tested on a set of 2.05M variants from 367 genes shared between VEPs **(a)**, and a set of 124,709 variants from 15 genes shared between VEPs and experimental MAVEs **(c). b**,**d**. Average MES and average AUROC show a strong correlation when tested on the large set of variants **(b)**, and the subset with MAVE data available **(d)**. All values are calculated on a per-gene level and then averaged across genes. Error bars denote standard errors of the mean.

In Fig. 2b, we compared MES to AUROC across all genes. Overall, the correlation is strong (Spearman’s ρ = 0.86), and CPT-1 ranks first by both metrics, but several methods show notable divergence. For example, SaProt ranks second in average MES, but sixth in average AUROC, while PHACT ranks fourth in AUROC but ninth in MES. In the extended analysis, EVE ranks first by MES but sixth by AUROC. These differences suggest that some VEPs are more effective at generating clinically relevant evidence than at binary variant classification.

We next considered MAVE data, comparing their experimentally determined variant effect scores to those from VEPs. In total, we calibrated 15 MAVE datasets for different human disease genes, covering 124,709 variants, and compared these to high-coverage VEPs (Fig. 2b-c). Intriguingly, MAVE data ranks first in terms of average MES, but performs markedly worse by AUROC, ranking lower than 7 of 12 VEPs. This suggests that functional measurements from MAVEs may be relatively better suited than VEPs to provide clinical evidence, even if their performance by AUROC is lower. However, MAVE data are now increasingly used for variant classification [22–24], and some variants in our calibration set are therefore likely to have been classified using these datasets, introducing circularity and potentially inflating the estimated evidence strength. For example, we recently highlighted an MSH2 variant that may have been misclassified as benign on the basis of MAVE data, leading to disagreement with VEPs [25]. It is also difficult to comprehensively identify pathogenic and benign variants for which MAVE data has contributed to classification, preventing us from reliably quantifying the extent of this circularity.

### MES Captures Clinical Evidence Yield Beyond Discrimination

To examine the relationship between conventional classification performance and clinical utility, we next compared AUROC to MES across the 367 disease genes, using CPT-1 because it performed best across both metrics. The relationship is strong (ρ = 0.89) and monotonic but non-linear, with MES increasing slowly across most of the AUROC range and rising sharply as AUROC approaches 1.0 (Fig. 3a). Accordingly, AUROC values above ∼0.95 almost always correspond to MES ≥ 2, whereas AUROC below ∼0.9 typically results in MES < 2. However, splitting MES into pathogenic and benign components (Fig. 3b) results in weaker correlations (ρ = 0.68 for pathogenic MES, 0.73 for benign MES). Thus, high AUROC does not necessarily translate into high clinical evidence in both directions, and often reflects strong evidence for either pathogenic or benign variants, but not both.

**Figure 3.**
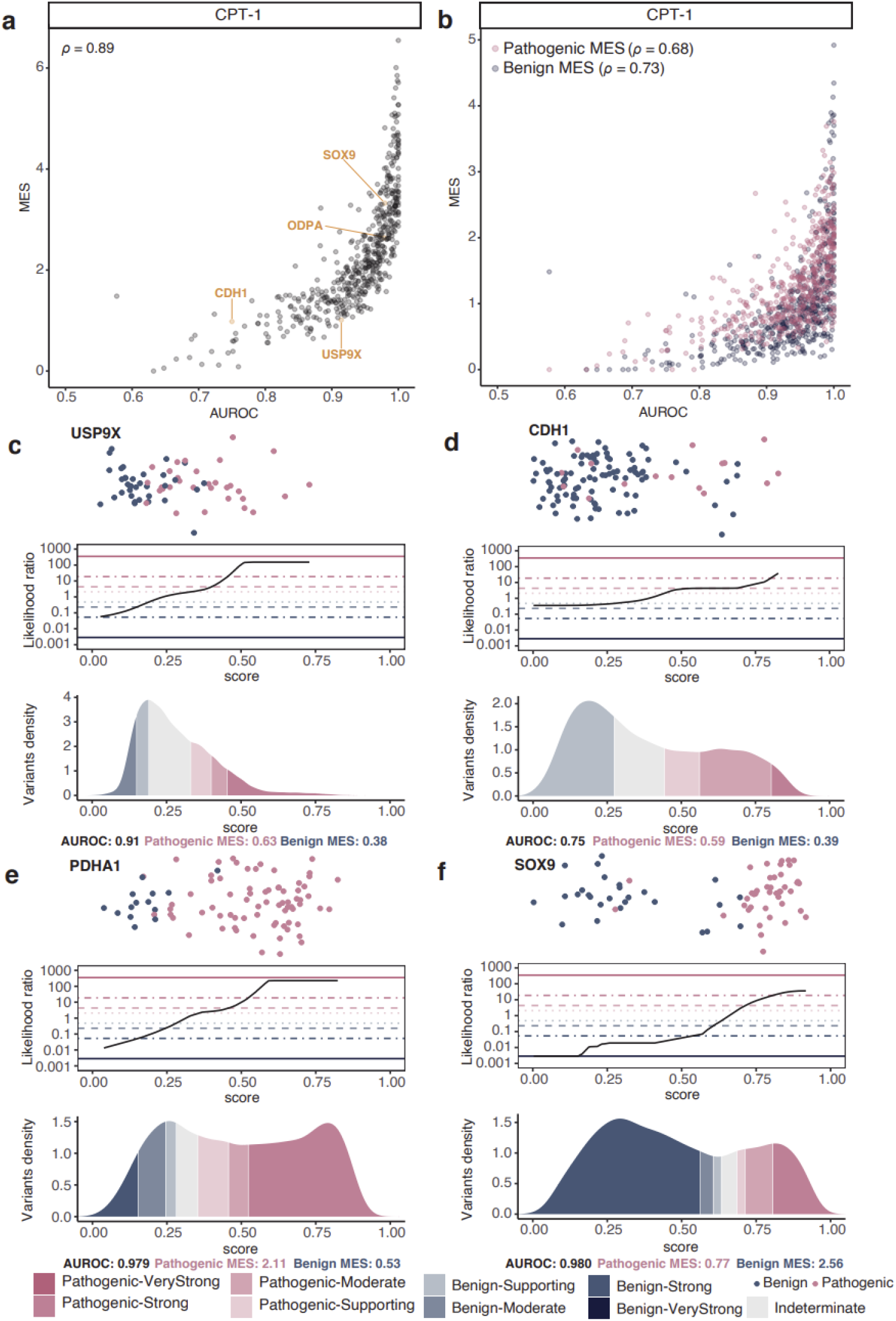
MES reveals gene-level clinical utility profiles beyond discrimination with AUROC. **a**. CPT-1 AUROC correlates well with MES. **b**. AUROC does not explain well the benign and pathogenic MES profiles using CPT-1 as an example. **c-f**. The CPT-1 score distribution of ClinVar benign and pathogenic variants (top panel), relationship between variant score and calibrated evidence thresholds (middle panel), and all variant score distribution coloured by evidence category (bottom panel) for USP9X, CDH1, PDHA1 and SOX9 as examples.

To further explore this discrepancy, in Fig. 3c-f we show the distributions of pathogenic and benign variants used for calibration, the *acmgscaler* likelihood ratio plots, and the distributions of calculated evidence levels across all variants for four selected genes. USP9X (Fig. 3c) has a relatively high AUROC of 0.91 but a low MES of 1.01. Examination of the distribution shows that, while there is generally good discrimination between pathogenic and benign variants, a large proportion of possible variants fall in the intermediate indeterminate region, thereby limiting the overall evidence yield. In comparison, CDH1 (Fig. 3d) has a similar MES of 0.98, but a much lower AUROC of 0.75. Here, although the overall discrimination between pathogenic and benign variants is relatively poor, the distribution of possible variants is slightly bimodal, with peaks in the *Benign-Supporting* and *Pathogenic-Moderate* regions. These examples show that MES depends not only on how well pathogenic and benign variants are separated, but also on how the full distribution of possible variants is positioned relative to the calibrated evidence thresholds.

PDHA1 (Fig. 3e) and SOX9 (Fig. 3f) have nearly identical AUROC values of 0.98, indicating excellent discrimination between pathogenic and benign variants. Both genes also show bimodal distributions of possible variants, with peaks on the pathogenic and benign sides. However, the balance between pathogenic and benign evidence differs markedly between them. In PDHA1, the pathogenic peak lies almost entirely within the strong-evidence region, resulting in a much higher pathogenic MES (2.11) than in SOX9, where only part of the pathogenic peak reaches strong evidence (pathogenic MES = 0.77). The opposite is true for benign evidence: in SOX9, the benign peak falls almost entirely within the strong-evidence region, giving a higher benign MES (2.56), whereas in PDHA1, only a smaller fraction of the benign peak does so (benign MES = 0.53).

### Mean Evidence Strength Reflects VUS Reclassification Utility

MES, as we have defined it, is calculated across all missense variants in a VEP or MAVE dataset. In practice, however, clinical utility depends on the ability to provide evidence for observed variants, particularly VUS, rather than on behaviour across all possible variants. In principle, MES could be calculated using only VUS, but this would introduce gene-specific ascertainment biases arising from differences in which variants have been submitted to ClinVar. To assess whether MES calculated across all variants nevertheless reflects practical reclassification utility, we compared average pathogenic and benign MES to the fraction of ClinVar VUS receiving at least moderate evidence across VEPs (Fig. 4a,b). The correlations were very strong (ρ = 0.97 for pathogenic MES and 0.86 for benign MES), and CPT-1 yielded the highest fraction of VUS receiving both pathogenic and benign evidence. Thus, MES calculated across all variants is highly reflective of the clinical evidence that a dataset can provide when applied to VUS.

**Figure 4.**
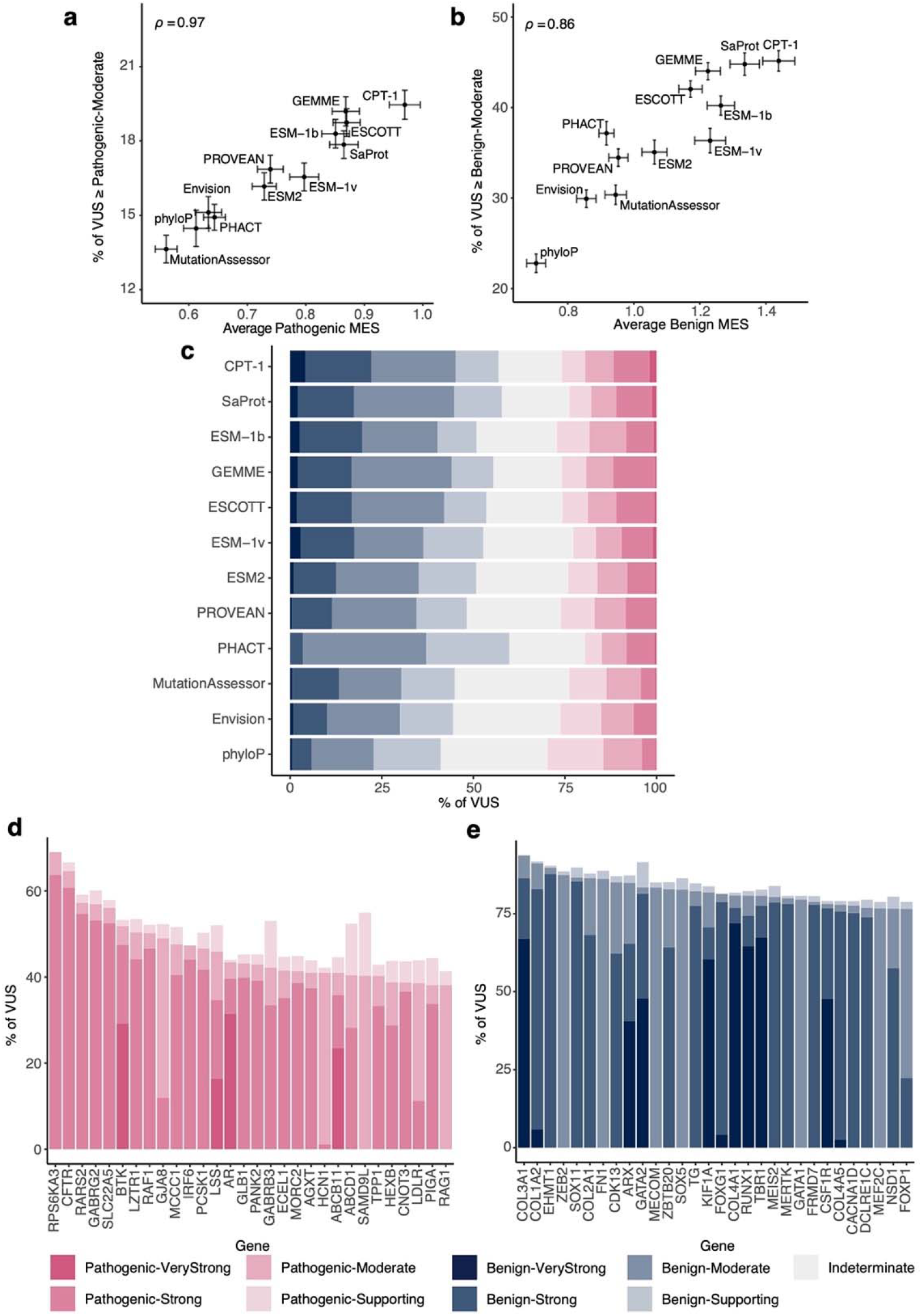
MES is reflective of clinical utility when applied to VUS. **a-b**. Average pathogenic **(a)** and benign **(b)** MES of VEPs correlate strongly with the fraction of VUS from ClinVar receiving at least moderate evidence from them. Error bars denote standard errors of the mean. **c**. Average fraction of VUS from each gene receiving different levels of evidence from each VEP. **d-e**. Top 30 genes where the highest fraction of VUS receive at least moderate pathogenic **(d)** and benign **(e)** evidence from CPT-1, showing the distribution of different evidence levels.

We next assessed the distribution of evidence assigned to ClinVar VUS across VEPs (Fig. 4c). All methods were able to assign strong pathogenic and benign evidence to a subset of variants, and several reached the very strong level. CPT-1 consistently provided evidence for the largest fraction of variants across most categories, although GEMME assigned strong pathogenic evidence to a slightly higher fraction, and some methods assigned supporting evidence more broadly. Overall, while there is variability in evidence profiles across methods, CPT-1 clearly provides the most evidence when applied to VUS. We also compiled the thresholds for 15 VEPs across all calibrated genes (Table S1).

We also assessed the sensitivity of our results to the choice of prior probability of pathogenicity. Throughout, we used the *acmgscaler* default of 0.1, as defined in the original Bayesian calibration framework [26]. We repeated the analyses using a lower prior of 0.0441, estimated from the prevalence of pathogenic variants among rare missense variants in Mendelian disease genes in gnomAD [7]. Pathogenic and benign MES calculated under the two priors were almost perfectly correlated, although absolute values were slightly lower with the lower prior (Fig. S2a,b). The distribution of evidence assigned to VUS was also similar overall, but with a higher fraction of variants remaining indeterminate and an almost complete loss of variants assigned very strong evidence (Fig. S2c). This is consistent with previous observations that lower priors lead to reduced evidence strengths [27].

Finally, to illustrate the potential utility of this approach for VUS interpretation, we ranked genes by the fraction of ClinVar VUS receiving pathogenic (Fig. 4d) and benign (Fig. 4e) evidence using CPT-1. Several genes show high coverage of VUS with pathogenic evidence, with a substantial proportion reaching at least moderate strength. For example, more than 60% of CFTR VUS are assigned strong pathogenic evidence. The corresponding patterns for benign evidence are even more pronounced, with many genes showing over three quarters of VUS receiving at least moderate benign evidence. Notably, several collagen genes, including COL3A1, show particularly strong enrichment for benign evidence, with more than 70% of VUS assigned very strong benign evidence.

## Discussion

Experimental and computational variant effect scores are increasingly discussed in terms of clinical utility, yet their performance is still usually summarised using classification metrics such as AUROC. Our results show that these metrics do not fully capture the extent to which a predictor or assay can contribute clinically interpretable evidence. Using a gene-level Bayesian calibration framework, we converted VEP and MAVE scores into standardised ACMG/AMP evidence categories and used these to define MES, a measure of evidence yield across a score distribution. Across 15 VEPs and 15 MAVE datasets, MES captured important properties that were only partly reflected by AUROC, particularly when pathogenic and benign evidence were considered separately. Calibration also exposed gene-specific differences in evidence profiles and was strongly associated with the proportion of ClinVar VUS receiving moderate or stronger evidence.

We found CPT-1 to provide the highest clinical evidence yield among the VEPs examined. This is consistent with our previous observations that CPT-1 shows the strongest agreement with experimental MAVE data and, among population-free VEPs, the best discrimination between pathogenic and benign variants [15]. We therefore recommend CPT-1 as the most suitable current choice for gene-level calibration for clinical evidence assignment. An alternative strategy could be to select the best-performing VEP separately for each gene, on the assumption that some predictors may perform better in specific gene contexts. In principle, this differs from the variant-level cherry-picking previously cautioned against [7], as selection would be performed at the gene level using an independent set of calibration variants, rather than choosing a predictor based on the evidence assigned to the specific variant under interpretation. However, such an approach could still inflate evidence through a winner’s curse-like effect, as some predictors will appear to perform best for particular genes partly by chance. We therefore favour the use of a single well-performing predictor over ad hoc per-gene selection, unless future work identifies principled reasons why particular VEPs should be preferred for specific genes.

Another important consideration is the choice of prior probability of pathogenicity. In principle, this prior should reflect the probability that a variant encountered in a clinical setting is pathogenic before considering the specific functional score or predictor output. In the original Bayesian reinterpretation of the ACMG framework, a value of 0.1 was proposed as a pragmatic approximation for panel-based testing [26]. More recently, efforts have been made to estimate this quantity empirically from the prevalence of pathogenic variants in reference population datasets such as gnomAD [11,27]. However, this quantity may diverge from the clinical prior, particularly in settings such as highly constrained dominant genes, where damaging variants are depleted from population datasets but may have a relatively high pre-test probability in an appropriate clinical context. An important advantage of gene-level calibration is that it enables the use of gene-specific priors, but further work is needed to determine how these should best be defined for clinical interpretation. Accordingly, our approach is designed to remain flexible to different prior assumptions.

The observation that MAVE datasets achieved higher MES than all VEPs, despite weaker discrimination by AUROC, is notable. This suggests that functional assays may provide stronger evidence across a broader range of variant effects. One possible explanation is that MAVEs capture intermediate-impact variants that contribute relatively little to discrimination-based metrics but can support graded evidence assignment after calibration. However, this interpretation is complicated by the potential for circularity, as MAVE data are increasingly incorporated into variant classification. We did not attempt to exclude such cases, and therefore some inflation of evidence strength cannot be ruled out. These results should therefore be interpreted cautiously, but they are consistent with the idea that experimental assays can complement computational predictors by contributing evidence that is not fully reflected in discrimination-based metrics.

Circularity is also likely to become an increasing concern for the use of VEPs in calibration. As computational predictors are more widely incorporated into clinical interpretation, variants in resources such as ClinVar will increasingly reflect evidence derived from VEPs [28]. This issue extends beyond predictors directly used in classification, since VEPs tend to be correlated with one another [29]. As a result, variants whose classification has been influenced by computational predictions may not provide independent evidence for calibration. Careful tracking of the evidence used in variant classification will therefore be important for the reliable application of calibration-based approaches.

We suggest that MES is most useful as a comparative metric and should be used alongside established measures such as AUROC or precision–recall when assessing the performance of VEPs and MAVE datasets on clinically relevant variants. Like AUROC, its value depends on the composition of the dataset and, in this case, the choice of prior probability of pathogenicity, so absolute values should be interpreted with caution. Instead, MES provides a complementary perspective on clinical evidence yield, which is particularly useful when reporting new MAVE datasets and when comparing computational and experimental approaches. Because MES can be calculated for both VEPs and MAVEs, it also provides a framework for considering how these sources of evidence might be combined. In principle, evidence from calibrated predictors and assays could be combined additively, but this assumes conditional independence, which is unlikely to hold given that both VEPs and MAVEs capture overlapping aspects of the same underlying functional constraints, as reflected in their strong correlation. How best to combine these sources of evidence therefore remains an open question. To facilitate its use, we have implemented MES as a standard output in the *acmgscaler* Colab notebook (https://github.com/badonyi/acmgscaler).

## Methods

### Data Sources

#### ClinVar Variant Sets

Calibration of variant scores requires variant subsets with known pathogenicity. All variant records from ClinVar were downloaded in May 2025 [1]. We defined our benign and pathogenic variant sets as all missense variants classified as benign or likely benign and pathogenic or likely pathogenic respectively. We excluded any variants with conflicting classifications.

#### Multiplexed Assays of Variant Effect Datasets

MAVE datasets used in this study were from MaveDB [30], ProteinGym [31], and our previous studies [15,25]. After mapping variant labels from ClinVar, we excluded any datasets with fewer than 10 pathogenic or benign variant labels as required by our gene-level calibration method [13].

#### Variant Effect Predictors Included

All VEPs previously defined as clinically trained were excluded from this study as they were exposed to clinical labels causing data circularity and potentially biasing calibration results [15]. We mapped ClinVar variant labels to predictions of each VEP and excluded any gene with fewer than 10 known pathogenic or benign variants as required by the *acmgscaler* package. After such filtering, we then excluded VEPs with low gene coverage (<400 genes). For fair comparison, the common set of variants covered by all remaining VEPs was determined and used for subsequent calibration. Finally, we included a set of 12 VEPs: CPT-1, ESM-1b [32], ESM-1v [33], ESM2 [34], ESCOTT [35], Envision [36], GEMME [37], MutationAssessor [38], PHACT [39], PROVEAN [40], SaProt [41], phyloP [42].

We also performed more lenient filtering (coverage > 350 genes) to include iGEMME [35], EVE and popEVE, as they are widely used predictors, but their inclusion in the main analysis would have reduced the common variant set by ∼50%.

### Gene-level Variant Effect Score Calibration

The R package *acmgscaler* was used to calibrate variant effect scores at the gene level [13] (Figure 1). Briefly, for any VEP or MAVE set, the distributions of known pathogenic and benign variants were modeled with separate Gaussian density kernels to estimate the likelihood of pathogenicity of variant scores normalised between 0 and 1. Using a prior probability of pathogenicity of 0.1 as previously defined [26], we calculated the posterior probability of pathogenicity of each variant given its likelihood ratio, which was then categorized into different evidence strengths (supporting, moderate, strong, very strong) using thresholds determined by confidence bounds of bootstrapping resamples. Thresholds were derived from the bootstrap confidence intervals of the likelihood ratio distributions. To maintain a conservative classification scheme, we applied the lower bound of the threshold interval when assigning benign evidence, and the upper bound when assigning pathogenic evidence.

### Mean Evidence Strength as a Scoring Metric

For a calibrated functional or predicted variant effect score set for any gene, mean evidence strength (MES) was calculated as the average absolute evidence point as previously defined (Supporting: 1, Moderate: 2, Strong: 4, Very-Strong: 8), received by all single amino substitutions in the score set [19] (Figure 1). Briefly, variants in scores set S with annotated pathogenic and benign label sets and can be assigned to pathogenic (*P*_*VSt*_, *P*_*St*_, *P*_*M*_, *P*_*Su*_) and benign (*B*_*VSt*_, *B*_*St*_, *B*_*M*_, *B*_*Su*_) subsets of different evidence strengths (Eq. 1). Pathogenic (*MES*_*P*_) and benign (*MES*_*B*_) MES for score set S is then defined as the weighted average of number of variants in each evidence subsets (Eq. 2,3). All variants calibrated to have indeterminate evidence receive an evidence point of 0. Together, the sum of pathogenic and benign MES of any calibrated score set equals its MES.

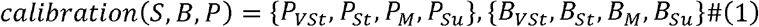

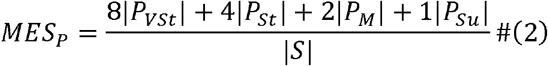

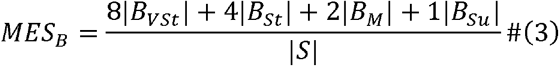

## Data Availability

Complete datasets associated with the analyses in this study are available at https://osf.io/2uxq9.

## Acknowledgements

This project was supported by funding to JAM from the European Research Council (ERC) under the European Union’s Horizon 2020 research and innovation programme (grant agreement No. 101001169) and by the Medical Research Council (MRC) Human Genetics Unit core grant (MC_UU_00035/9).

## Supplementary Figures

**Figure S1.**
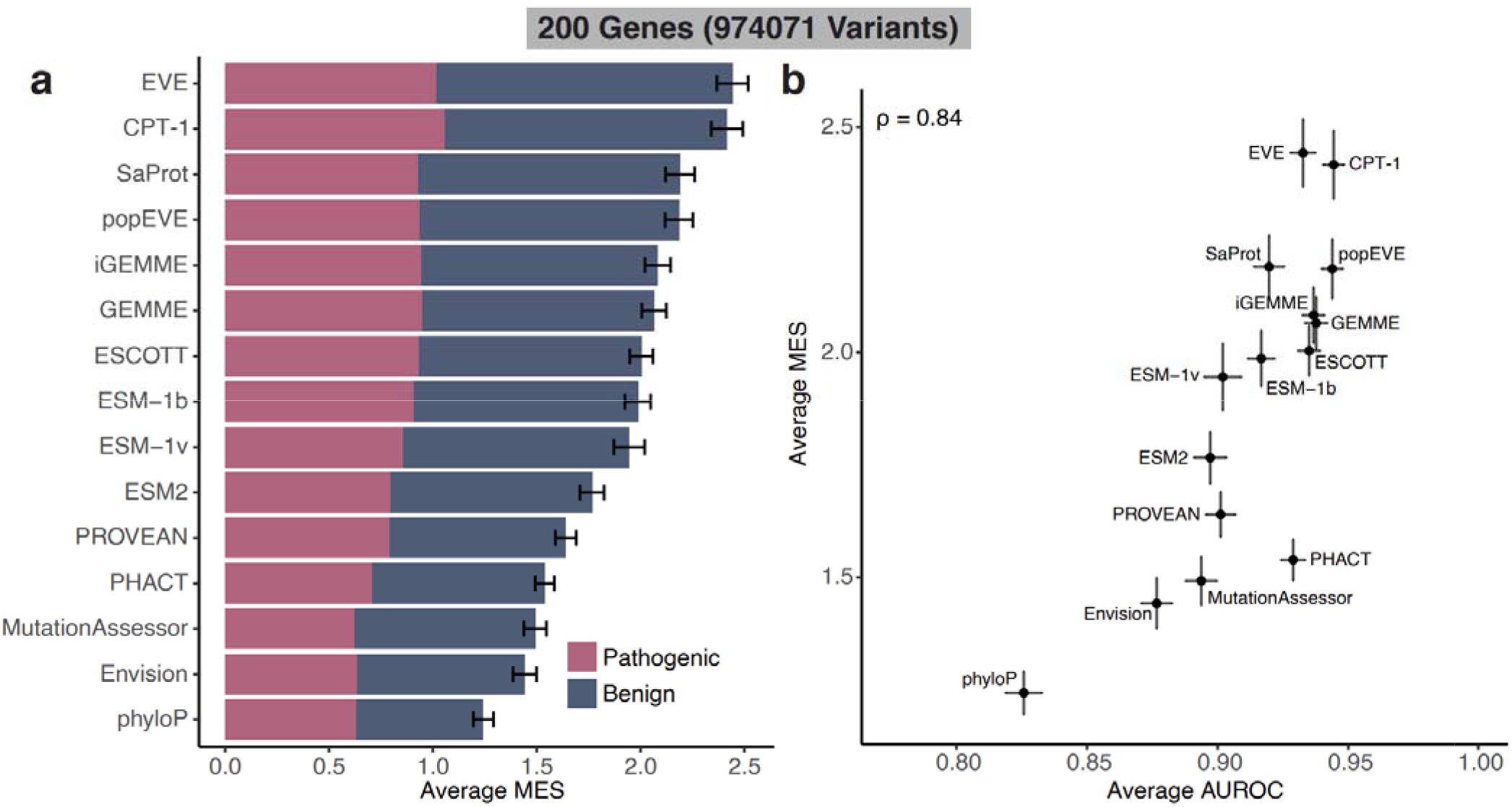
Measuring clinical utility of an expanded set of VEPs using the MES metric. Same as Fig. 2a-b, but using a smaller set of 974,071 variants from 200 genes, to enable the inclusion of further VEPs (EVE, popEVE and iGEMME).

**Figure S2.**
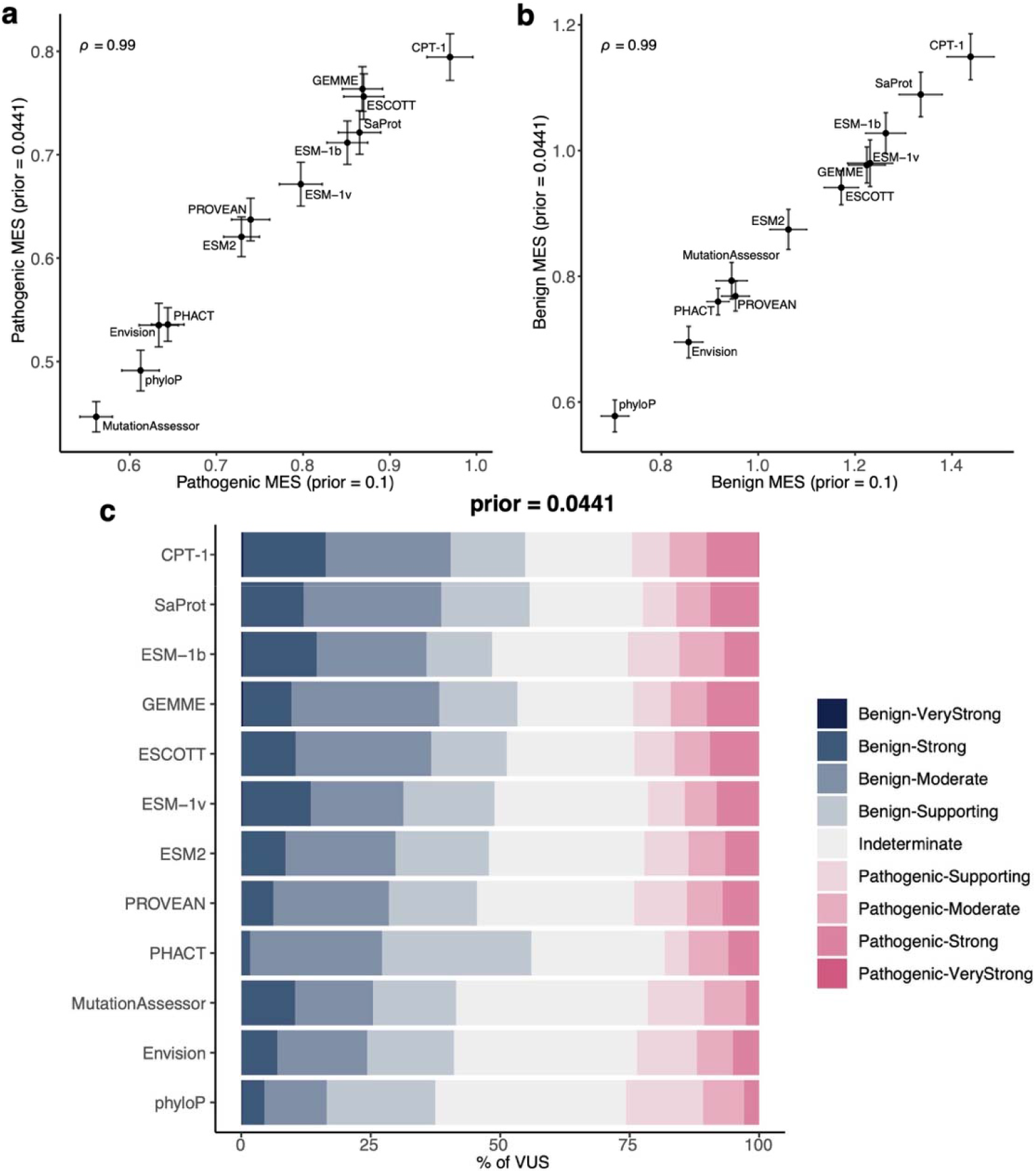
Influence of the prior probability of pathogenicity on MES and VUS reclassification. **a-b**. Comparison of pathogenic **(a)** and benign **(b)** MES calculated with the default prior of 0.1 and the stricter value of 0.0441. Error bars denote standard errors of the mean. **c**. Same as Fig. 4c but using the lower prior.

